# Structural analysis of OCT4 Binding to Human LIN28B Nucleosomes

**DOI:** 10.1101/2025.09.12.675874

**Authors:** Kalyan K Sinha, Mario Halic

## Abstract

Structural studies of nucleosomes most commonly involve histones from *Xenopus* species or humans. Yet, the effect of subtle differences in the amino acid sequences of these histones on key aspects of structure, such as nucleosome assembly, DNA positioning, and transcription factor binding remains unclear. Here, we show that histones from both species can efficiently assemble on the human *LIN28B* DNA sequence. Using cryogenic electron microscopy we demonstrate that the pioneer transcription factor OCT4 engages with LIN28B nucleosomes assembled with human histones in the same manner as observed in our previous work in which the nucleosomes were assembled with *Xenopus* histones.

## Introduction

Pioneer transcription factors (TFs) are a specialized class of TFs characterized by their unique ability to access DNA within compact, transcriptionally inactive chromatin^1,2^. Pioneer TFs can cooperate with one another to regulate chromatin accessibility and influence cell fate decisions. For example, the induction of pluripotency is mediated by the cooperative action of pioneer TFs, such as OCT4, SOX2, KLF4, and cMYC^3^ Despite their central roles in development and cellular reprogramming, the molecular mechanisms by which pioneer TFs engage with nucleosomes and act cooperatively remain incompletely understood.

Several mechanisms have been proposed to explain how pioneer TFs bind to nucleosomes. We recently reported the structure of full-length OCT4 bound to nucleosomes assembled with a 182-bp DNA sequence from the regulatory region of the human *LIN28B* locus^4^. We demonstrated that OCT4 binding to the linker DNA region (i.e., OCT4 binding site 1 [OBS1]) stabilizes DNA positioning on the nucleosome, thereby increasing the accessibility of internal binding sites to OCT4 and SOX2. We also found that acetylation of the histone H3 residue K27 (H3K27ac) further promotes DNA repositioning and enhances the binding of OCT4 to its internal nucleosomal sites, OBS2 and OBS3. Finally, we determined the structure of OCT4 bound to nucleosomes assembled with the DNA sequence from the *nMATN1* locus, which showed that OCT4 engages with the nMATN1 nucleosomes in a manner similar to its binding with the LIN28B nucleosomes. This finding suggests that linker DNA binding is a more general mode of OCT4 interaction with nucleosomes.

The nucleosomes used for structure determination in our published study contained histones from *Xenopus laevis*, which display a high degree of sequence similarity with the corresponding human histones. Yet, there are some differences in the amino acid sequences between the two species in histones H2A and H2B. Although these differences do not affect the nucleosome structure, they may change the engagement of sequence-specific TFs with nucleosomes.

In the present study, we investigated whether *Xenopus* or human histones affect DNA positioning and OCT4 engagement with LIN28B nucleosomes. We show that the assembly of the nucleosomes and their positioning on DNA did not change between reconstitutions that use *Xenopus* or human histones. We then used cryogenic electron microscopy (cryo-EM) to solve the structure of the OCT4–LIN28B nucleosome complex assembled with human histones. Our findings demonstrate that OCT4 binds nucleosomes containing *LIN28B* DNA and *Xenopus* or human histones in the same way.

## Results

### Nucleosome assembly and positioning are unaffected by histone origin or assembly method

The amino acid sequences of *Xenopus* histones and human histones show a high degree of evolutionary conservation, with 15 of 17 (88%) differences confined to histones H2A and H2B (**Supplementary Fig. 1, 2**). To test whether such differences affect the mode of engagement of OCT4 with nucleosomes, we assembled OCT4 with LIN28B nucleosomes containing human and *Xenopus* histones.

To assemble nucleosomes, we used a double-bag dialysis method^5-7^ (**Fig. 1A**) that provides slow salt dilution, similar to the salt-gradient dialysis method described by Luger et al^8^. We assembled nucleosomes by using *LIN28B* DNA and either human or *Xenopus* histones. Native gel electrophoresis analysis showed that histone octamers from both species efficiently assembled into nucleosomes (**Fig. 1B**). Additionally, SDS gel electrophoresis showed that the band intensities of histones are indicative of nucleosome assembly (**Fig. 1C**). These data confirmed that the double-bag dialysis method supports robust assembly of nucleosomes with *Xenopus* or human histone octamers.

**Fig. 1.**
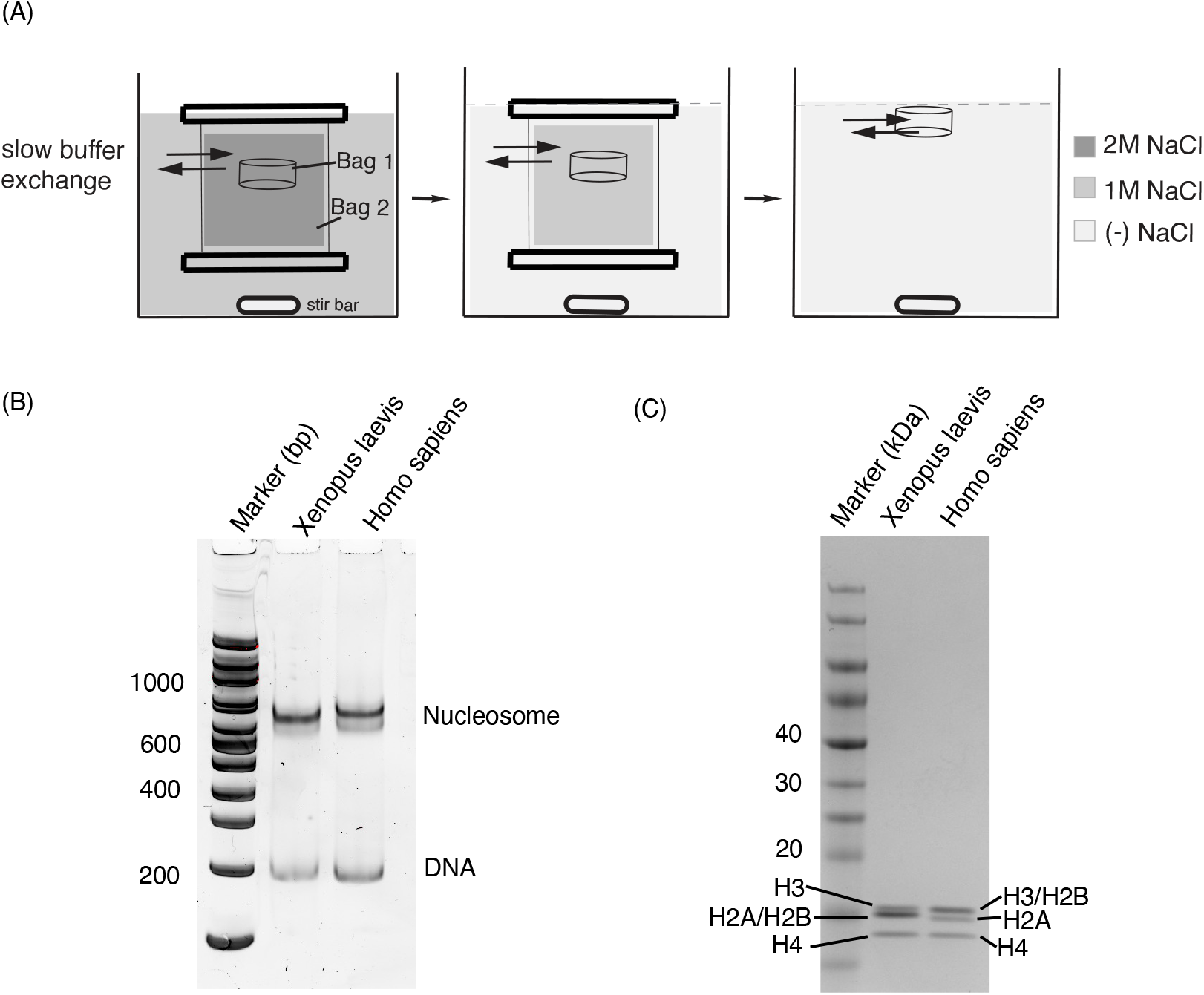
Assembly of LIN28B nucleosomes. **(A)** Schematic for double-bag dialysis method used for nucleosome assembly. **(B)** Native PAGE analysis of the 182-bp LIN28B nucleosomes assembled with *Xenopus* or human histone octamers via the double-bag dialysis method. **(C)** SDS PAGE analysis of LIN28B nucleosomes assembled with *Xenopus* or human histone octamers by using the double-bag dialysis method.

In our previous study, we observed LIN28B nucleosomes in multiple positions. To determine whether prolonged heating at physiological temperature can drive the multiple nucleosome positions to a single position on the *LIN28B* DNA, we heated the assembled nucleosomes at 37 °C for 5 h. If heating causes nucleosome repositioning, then the repositioned products should appear on a native gel, either as a new band(s) or as an increased fraction of the band below the nucleosome band. Instead, our native PAGE analysis (**Fig. 2A**) showed that the nucleosomes did not undergo repositioning after heating. To confirm our observation, we used an orthogonal approach, MNase-seq (micrococcal nuclease sequencing) analysis of nucleosome samples, with or without heating, and found that heating did not change the position (**Fig. 2B**). Together, these results indicated that the assembled LIN28B nucleosomes inherently occupy multiple positions, even at physiological temperature.

**Fig. 2.**
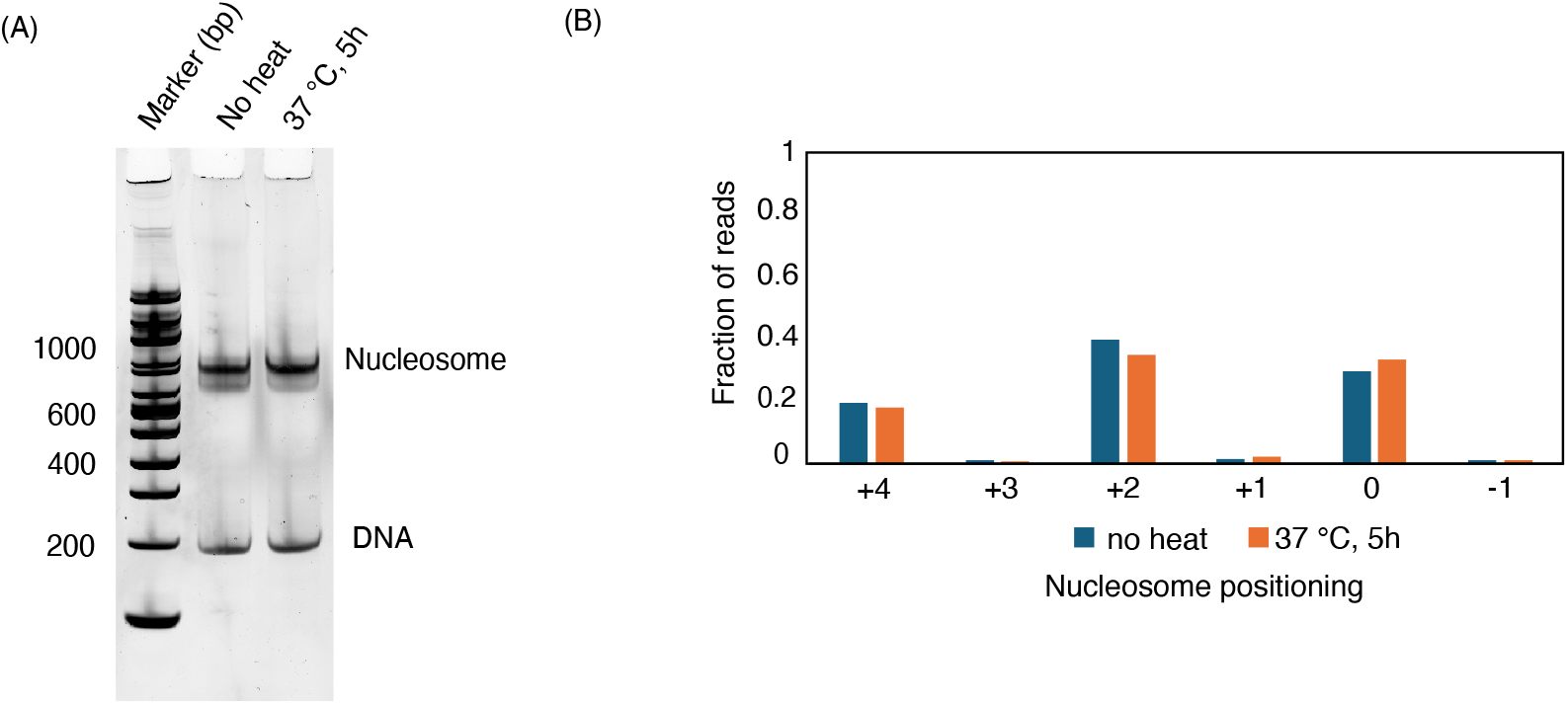
Effect of heating on the positioning of LIN28B nucleosomes containing *Xenopus* histones. **(A)** Native PAGE analysis of nucleosomes with or without heating at 37 ° C for 5 h. The second lower band is unaffected. **(B)** MNase-seq analysis of nucleosome positioning on *LIN28B* DNA.

### Structure of the OCT4–LIN28B nucleosome complex assembled with human histones

Next we assembled an OCT4–LIN28B nucleosome complex by adding OCT4 to pre-assembled human nucleosomes. The structure of the OCT4–LIN28B nucleosome complex was then determined from ∼15,000 particles by using cryo-EM at 6 Å resolution. The structure revealed that both DNA-binding domains of OCT4 (i.e., POU_S_ and POU_HD_) engaged linker DNA near the entry/exit site of the nucleosome (**Fig. 3A**), at the same location described by Sinha et al.^4^. This result indicates that histone sequence variations between human histones and *Xenopus* histones do not affect the mode of OCT4 engagement with the LIN28B nucleosome. Indeed, our previous model of OCT4– LIN28B nucleosome (PDB 8g8g) fit well into the map of OCT4 bound to human LIN28B nucleosome (**Fig. 3B**). Thus, the structure of the OCT4–LIN28B nucleosome remains the same.

**Fig. 3.**
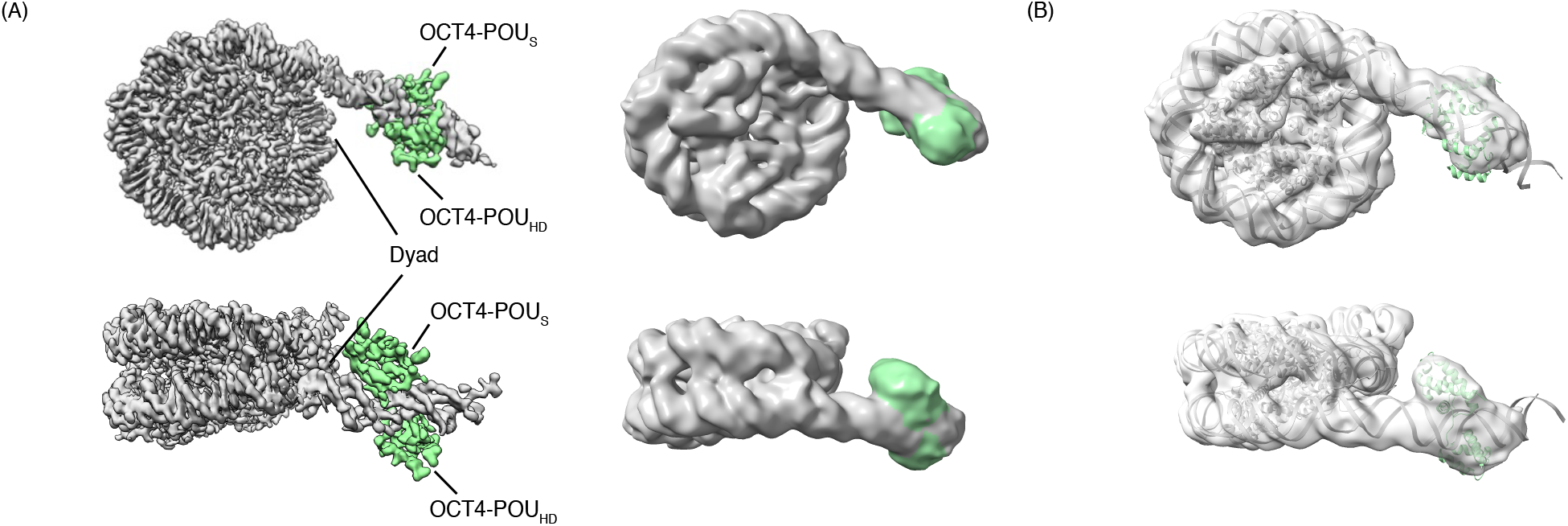
**(A)** Cryo-EM structure of the LIN28B nucleosome assembled with human histones bound to OCT4 (green); shown in top and side views. Left, OCT4 bound to the LIN28B nucleosome with *Xenopus* histones. Right, OCT4 bound to the LIN28B nucleosome with human histones. POU_S_, POU-specificity domain; POU_HD_, POU homeodomain. **(B)** Fitting of the model of the OCT4–LIN28B nucleosome assembled with *Xenopus* histones (PDB 8G8G) on the density map of the OCT4–LIN28B nucleosome assembled with human histones (this work, EMDB EMD-72371). The overlay was performed using rigid body fitting in ChimeraX.

## Discussion

Here, we showed that the OCT4 TF binds the same site (i.e., OBS1) on LIN28B nucleosomes, regardless of whether it is assembled with *Xenopus* histones or human histones.

In previous work, we showed that *LIN28B* DNA occupies multiple positions on nucleosomes and the binding of OCT4 to nucleosomes preferentially selects the position of the DNA seen in the cryo-EM structure. In the present study, we have demonstrated that the LIN28B DNA adopts multiple positions on the histone octamer, in the context of both *Xenopus* and human histones, suggesting that the sequence differences between the histones have minimal effect on DNA positioning, and variable nucleosomal DNA positioning could be a feature of nonpositioning native sequences. Our observations are in agreement with the chemical mapping data that showed *LIN28B* DNA can adopt multiple positions on nucleosomes^9^. Consistent with this finding, in vivo positioning data have shown a much broader peak (∼200-250 bp) of nucleosome occupancy at the *LIN28B* locus (as reported in GSE49140)^10^.

The mode of engagement of OCT4 with LIN28B and nMATN1 nucleosomes in this study and our previous study, shows that OCT4 binding stabilizes one position among the multiple DNA positions on the nucleosome, thereby promoting cooperative TF binding on nucleosomal DNA. Such behavior has recently been observed for other TFs, suggesting that this is a more widely used mechanism of pioneer TFs than previously thought^11^.

## Methods

### Protein expression and purification

Human histones were purchased from The Histone Source (Colorado State University). *Xenopus laevis* histones were overexpressed in *Escherichia coli* BL21(DE3) pLysS cells and purified from inclusion bodies, as previously described^12,13^.

Cells were maintained in culture in LB medium at 37 °C and induced with 1 mM IPTG, when the optical density at 600 nm (OD_600_) reached 0.6. After 3 h of expression, the cells were harvested by centrifugation, resuspended in lysis buffer (50 mM Tris-HCl, pH 7.5; 150 mM NaCl; 1 mM EDTA; 1 mM DTT; and 0.1 mM PMSF), and frozen. The frozen cells were later thawed and lysed by sonication. Inclusion bodies were collected by centrifugation at 5,000 rpm for 20 min at 4 °C.

The resulting pellet was washed three times with a lysis buffer containing 1% Triton X-100, followed by two additional washes with lysis buffer lacking Triton X-100. Individual histone proteins were extracted from the purified inclusion bodies by incubation in a denaturing buffer (50 mM Tris, pH 7.5; 2 M NaCl; 6 M guanidine hydrochloride; and 1 mM DTT) overnight at room temperature. Insoluble debris was removed by centrifugation at 13000 rpm 10 min.

Equimolar amounts of histone pairs (H2A–H2B and H3–H4) were mixed and subjected to two rounds of dialysis in 1 L refolding buffer (25 mM HEPES-NaOH, pH 7.5; 2 M NaCl; and 1 mM DTT) at 4 °C. Precipitated material was removed by centrifugation at 13,000 rpm for 20 min at 4 °C. The soluble histone pairs were further purified via batch-mode cation-exchange chromatography using SP Sepharose Fast Flow resin. Samples were diluted fourfold with salt-free buffer (25 mM HEPES-NaOH, pH 7.5; 1 mM DTT) and incubated with the resin for 30 min. The resin was washed extensively with 500 mM NaCl buffer (25 mM HEPES-NaOH, pH 7.5; 500 mM NaCl; and 1 mM DTT) and then transferred to a disposable column. Proteins were eluted with high-salt buffer (25 mM HEPES-NaOH, pH 7.5; 2 M NaCl; and 1 mM DTT), concentrated, and subjected to size-exclusion chromatography on a Superdex S200 column (GE Healthcare) equilibrated in the same buffer. Fractions containing pure histone proteins were pooled, concentrated, and flash-frozen for storage.

For cryo-EM and other assays, His-tagged OCT4 (∼39 kDa) was expressed from a pET28 vector and purified under denaturing conditions from inclusion bodies by using Talon affinity resin. Refolding was carried out by stepwise dialysis: the first dialysis was performed overnight in a buffer containing 2 M urea, 50 mM HEPES (pH 7.5), 250 mM NaCl, and 2 mM DTT. This was followed by two successive 1-h dialyses in a buffer containing 50 mM HEPES (pH 7.5), 100 mM NaCl, and 1 mM DTT.

### Histone octamer assembly and purification

Histone octamers were purified using the standard protocol^12,13^. Briefly, a 2.5-fold molar excess of the H2A–H2B dimer was mixed with the H3–H4 tetramer in refolding buffer (25 mM HEPES, pH 7.5; 2 M NaCl; and 1 mM DTT). The mixture was incubated overnight at 4 °C to allow octamer assembly. Excess H2A–H2B dimers were removed by size-exclusion chromatography using a Superdex S200 Increase 10/300 GL column on an ÄKTA FPLC system. Eluted fractions were analyzed by SDS–PAGE, pooled, concentrated, and used for nucleosome assembly.

### Nucleosome assembly

Nucleosomes were assembled using a double-bag dialysis method as previously described^5-7^. The histone octamer and nucleosomal DNA fragment were mixed in equimolar ratios in a buffer containing 50 mM HEPES (pH 7.5), 2 M NaCl, and 2 mM DTT. The mixture was placed into a dialysis button made with a membrane with a cut-off of 3.5 kDa. The dialysis button was placed inside a dialysis bag (6-to 8-kDa cut-off membrane) filled with 50 mL buffer containing 25 mM HEPES (pH 7.5), 2 M NaCl, and 1 mM DTT. The dialysis bag was immersed into 1 L buffer containing 25 mM HEPES (pH 7.5), 1 M NaCl, and 1 mM DTT, and dialyzed overnight at 4 °C. The next day, the buffer was changed to 1 L buffer containing 25 mM HEPES (pH 7.5) and 1 mM DTT, and dialysis was continued for 6–8 h. In the last step, the dialysis button was removed from the dialysis bag and dialyzed overnight into a fresh buffer without any salt (50 mM HEPES [pH 7.5] and 1 mM DTT). The nucleosome assemblies were assessed on a 6% native PAGE gel using SYBR gold staining.

### MNase-seq

LIN28B nucleosomes samples were digested by MNase (NEB) for 5 min at 25 °C (50 mM HEPES (pH 7.5), 1 mM DTT, 50 mM KCl, 5 mM CaCl_2_). MNase digestion was terminated by 50 mM EDTA. Cleaved nucleosomes were subjected to phenol/chloroform extraction followed by ethanol precipitation of nuclesomal DNA and used for library preparation. The sequencing library was prepared using the NEBNext Ultra II DNA Library Prep Kit following the manufacturer’s manual. The amplification of the library for Illumina sequencing was performed by PCR using NEBNext Multiplex Oligos for the Illumina kit. The sequencing was pair ended with 100-bp length. Paired reads were merged and filtered by the length of reads between 144 bp and 146 bp and mapped to the LIN28B sequence with Qiagen CLC genomics Workbench 20 software.

### Assembly of the OCT4–LIN28B complex for cryo-EM imaging

To assemble the nucleosome–OCT4 complex, we mixed pre-assembled LIN28B nucleosomes with OCT4 in a 1:5 molar ratio. The sample was incubated at 25 °C for 10 min and then transferred to ice for further processing.

### Cryo-EM grid preparation and data collection

The nucleosome–OCT4 complex (3 µL) was applied to freshly glow-discharged Quantifoil R1.2/1.3 300 mesh holey carbon grids. The humidity in the chamber was 95%, and the Vitrobot chamber temperature was set to 10 °C. After 5 s of blotting time, grids were plunge-frozen in liquid ethane by using an FEI Vitrobot automatic plunge freezer.

On a Talos Arctica electron microscope set at 200 kV with a Gatan Summit K3 electron detector, 436 micrographs were recorded using EPU software. Image pixel size was 1.06 Å per pixel on the object scale. Data were collected in a de-focus range of 7,000– 30,000 Å, with a total exposure of 60 e− Å^−2^. Sixty frames were collected and aligned with the Patch Motion Correction in the cryoSPARC software^14^. The contrast transfer function (CTF) parameters were determined using the Patch CTF Estimation job in cryoSPARC software^14^.

Initial templates were generated using the blob picker job in cryoSPARC, and ∼400,000 particles were picked using the templates. Based on the 2-dimensional (2D) class averages, several thousand junk particles were discarded from this set, resulting in ∼74,000 particles for ab initio generation of the initial 3D model. The initial model was refined to an ∼4 Å 3D map by using the Homogeneous Refinement job in cryoSPARC. To resolve the bound and unbound species, the map was classified into four 3D classes: class I and II had unbound nucleosomes, but class III and IV showed clear density for bound OCT4. The classes were further refined using the Heterogenous Refinement job in cryoSPARC. The output from the heterogenous refinement was used for reconstructing the final 3D map with 6 Å resolution using the gold standard FSC method in cryoSPARC.^14^

The previously published model, PBD 8G8G,^4^ was a rigid body fitted with the electron density map of OCT4–LIN28B nucleosome complex in ChimeraX. The cryo-EM maps were visualized using Chimera.

## Acknowledgements

We thank Cryo-EM Center members at St. Jude Children’s Research Hospital for support with grid screening and data collection; I. Chen, S. Bilokapic, and Angela McArthur for critical reading and comments; Janet Patridge for human histones; Genome Sequencing Facility at the Hartwell Center at St. Jude for Illumina sequencing of MNase-Seq libraries.

## Author contributions

K.K.S. and M.H. designed the experiments. K.K.S. performed the cryo-EM and biochemical experiments. K.K.S. and M.H. wrote the paper.

## Data availability statement

The EM density map has been deposited in the Electron Microscopy Data Bank under the following accession code: EMD-72371.

## Additional Information

The authors declare no competing interests.

## Supplementary Information

**Supplementary Figure 1.**
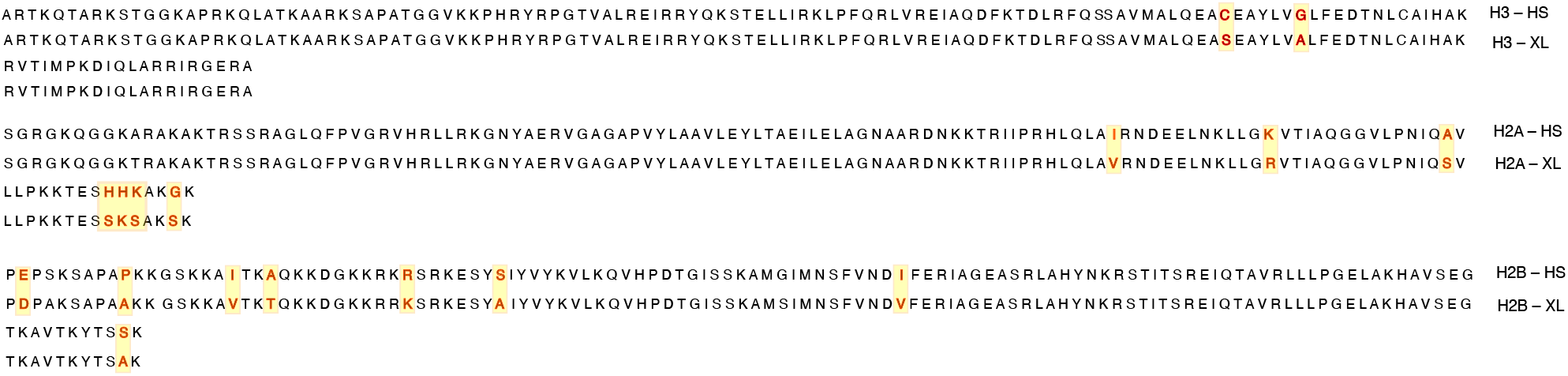
Sequence alignment showing differences between human histones and *Xenopus* histones (highlighted in yellow).

**Supplementary Figure 2.**
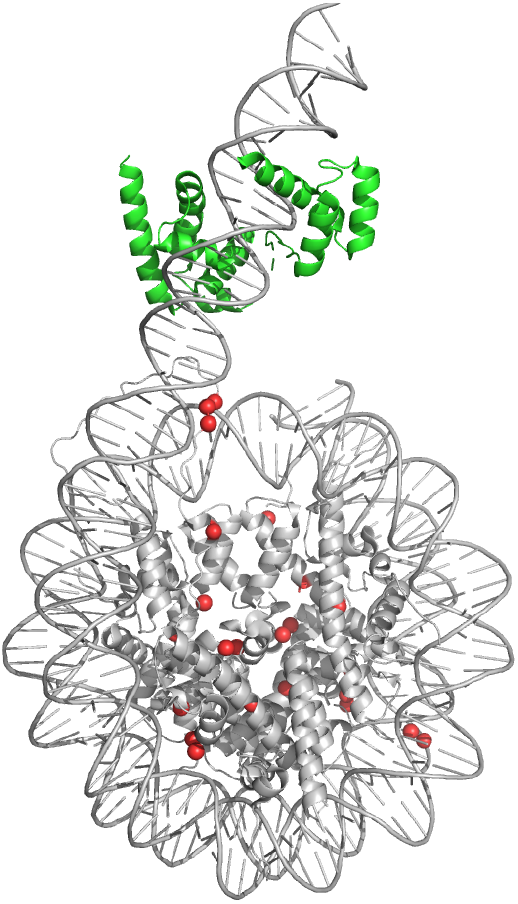
The locations of the amino acid residues that differ between *Xenopus* histones and human histones are mapped as red spheres on the structure of the OCT4-bound LIN28B nucleosomes assembled with *Xenopus* histones.

